# Monoaminergic changes are associated with impaired neurogenesis along the longitudinal axis of the hippocampus in ageing and dementia

**DOI:** 10.64898/2026.11.03.764096

**Authors:** Kathryn J Mathews, Amr H Abdeen, David Mor, Veronica Cottam, Marshall Axel Dalton, John B Kwok, Kay L Double

**Affiliations:** Brain and Mind Centre, The University of Sydney, Sydney, NSW, Australia; School of Medical Sciences (Neuroscience), Faculty of Medicine & Health, The University of Sydney, Sydney, NSW, Australia; School of Psychology, Faculty of Science, The University of Sydney, Sydney, New South Wales, Australia, 205

**Keywords:** Parkinson disease dementia, ageing, hippocampus, neurogenesis, neuronal maturation

## Abstract

Neurogenesis persists in the adult human hippocampus and is thought to support memory and emotional regulation. Both normal ageing and Parkinson disease dementia (PDD) are associated with reduced hippocampal neurogenesis, but why this occurs and how this might change across the across the hippocampal axis remains unclear. Here, we quantified the expression of genes regulating neurogenesis from stem cell production to neuron maturation and examined their relationship with monoaminergic activity throughout the post-mortem hippocampus in young and aged individuals, as well as in individuals with PDD. *DCX* and *MKI67* expression were markedly reduced in PDD cases throughout the hippocampus, while *GFAP* and *GFAPδ* were elevated in aged control individuals but not in PDD. Dopamine and serotonin levels were unchanged but reduced levels of the dopamine metabolite DOPAC, and increased levels of the serotonin metabolite 5-HIAA, were present across the hippocampal axis in PDD. Hierarchical regression analysis identified DOPAC and 5-HIAA as significant predictors of *DCX* and *MKI67* expression independent of diagnostic group. Coupling between stem cell markers and proliferative output was preserved in the hippocampus of young controls but was absent in aged controls and in PDD. These data suggest changes in monoaminergic systems are linked to stage-specific deficits in neuronal proliferation and maturation, identifying a potential mechanistic bridge to cognitive decline in PDD.

## INTRODUCTION

Adult hippocampal neurogenesis provides a continuing source of new granule neurons in the dentate gyrus and is thought to support circuit plasticity, pattern separation, and aspects of learning, mood, and stress regulation in both rodents and humans (Balu & Lucki, 2009). The extent and regulation of neurogenesis across the lifespan have been debated in humans, but current data suggest new neurons continue to be generated into old age, albeit at lower levels than in younger adults (Boldrini et al., 2018; Moreno-Jiménez, Terreros-Roncal, Flor-García, Rábano, & Llorens-Martín, 2021; Sorrells et al., 2018; Tobin et al., 2019). We previously showed that ageing is accompanied by reductions in markers of proliferating precursors and immature neurons in the hippocampus, while stem cell-associated markers remain relatively stable, pointing to an age-associated change in cellular maturation rather than the loss of stem cells (Mathews et al., 2017).

The hippocampus is anatomically and functionally heterogeneous along its longitudinal axis. Human imaging and rodent studies indicate that the anterior regions are closely linked to emotional processing and stress regulation, whereas the posterior regions are more engaged in spatial and episodic memory (Przeździk, Faber, Fernández, Beckmann, & Haak, 2019; Strange, Witter, Lein, & Moser, 2014). Neurogenesis rates and the composition of the progenitor pool vary along the dorsoventral hippocampal axis in rodents, with several studies reporting higher levels of cell proliferation and immature neurons in the tail of the hippocampus, preferentially supporting cognitive functions (Kozareva, Cryan, & Nolan, 2019; Poppenk, Evensmoen, Moscovitch, & Nadel, 2013; Strange et al., 2014). At the same time, the heterogeneous distribution of dopaminergic and serotonergic projections to the hippocampus suggests neuromodulatory systems may influence the number of neural progenitors throughout the hippocampus in rodents (Berg, Belnoue, Song, & Simon, 2013; Strange et al., 2014). Whether comparable regional gradients exist in the human hippocampus, and how they are affected by ageing and disease, remains largely unexplored.

Serotonin (5-HT) signalling may increase progenitor proliferation and promotes survival of newborn neurons, while dopamine (DA) and other neuromodulators can influence progenitor activity and integration in a state-dependent manner (Berg et al., 2013; Dranovsky & Hen, 2006). The distribution of dopaminergic and serotonergic projections to the hippocampus are topographically organised along the longitudinal axis in rodents (Gasbarri, Sulli, & Packard, 1997; Tsetsenis, Broussard, & Dani, 2023a), raising the possibility that regional variation in monoaminergic tone contributes to spatial differences in neurogenic activity. Critically, both monoaminergic systems are vulnerable to age-related changes and are further disrupted in neurodegenerative disease, suggesting that monoaminergic changes could contribute to impaired neurogenesis in humans (Kempermann et al., 2018; Shohayeb, Diab, Ahmed, & Ng, 2018). Parkinson disease dementia (PDD) provides a physiological context to examine this relationship. It is a neurodegenerative condition affecting the hippocampus in which both dopaminergic and serotonergic pathways are progressively compromised (Halliday, Leverenz, Schneider, & Adler, 2014; Perry et al., 1993) and in which cognitive decline, including memory impairment, is a prominent clinical feature (Aarsland et al., 2021).

Despite these links between ageing, monoaminergic alterations and cognitive decline, it is unknown if differences in dopaminergic and serotoninergic activity along the human hippocampal axis are associated with successive stages of the neurogenic lineage, nor if such relationships are altered in disease. Here, we used human post-mortem hippocampal tissue from young and aged healthy adults, as well as PDD cases, to investigate regional variations in the expression of neurogenesis-related genes and their relationship with dopamine and serotonin levels, and their metabolites.

## MATERIALS AND METHODS

### Human post-mortem brain tissue

Human post-mortem brain tissues from young adult individuals (age: 18–41; *n* = 12), aged adult individuals (age: 55–97; *n* = 9), and PDD cases (age: 60–90; *n* = 9) were obtained from the National Institute of Health Neurobiobank (**Table 1**). Fresh-frozen tissue samples were provided from the head, body, and tail of the hippocampus. Hippocampal body was defined as the region at the coronal level of the lateral geniculate nucleus, while the head and tail hippocampus were defined as 1–1.5 cm anterior and posterior to the body hippocampus, respectively (**Fig. 1**). Tissue samples were supplied as either hippocampal tissue punches or as a coronal block containing the hippocampus. Tissue punches were pulverised on dry ice using a mortar and pestle and distributed randomly into centrifuge tubes with tissue weight between 50–100 mg. For coronal blocks, 50 µm sections were cut using a freezing microtome and distributed sequentially into centrifuge tubes with tissue weight between 50–100 mg.

**Figure 1.**
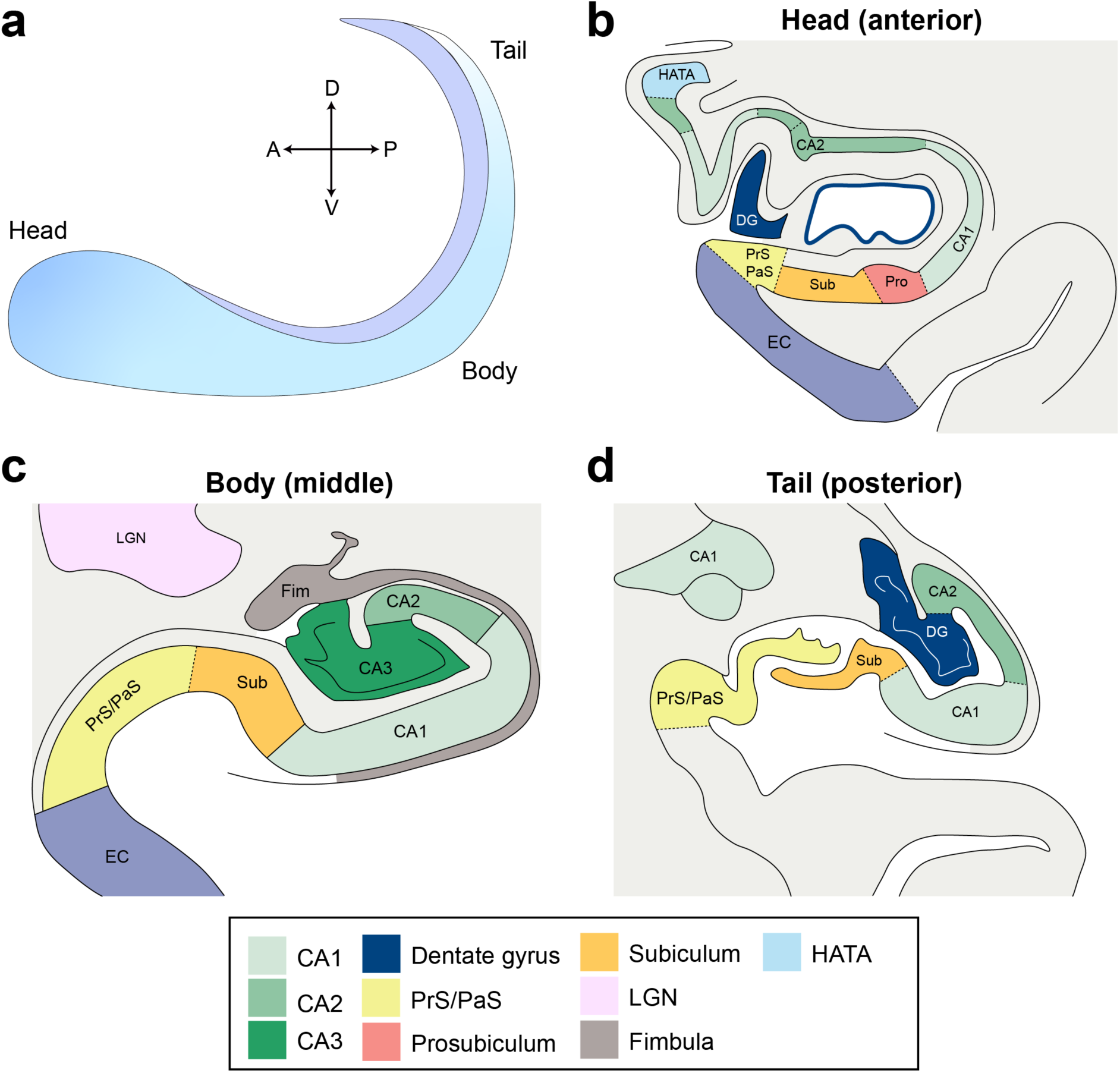
Regional organisation of the human hippocampus along its longitudinal axis. **(a)** Schematic of the hippocampal long axis showing the anatomical division into head, body, and tail. The body was defined at the coronal level of the lateral geniculate nucleus, and the head and tail were defined as 1 to 1.5 cm anterior and posterior to this reference level, respectively. **(b–d)** Representative coronal schematics illustrating the subregional architecture of the hippocampus at the head (anterior), body (middle), and tail (posterior) levels. These sections highlight the changing topography and relative boundaries of major hippocampal and parahippocampal subfields across the longitudinal axis. *Abbreviations: A, anterior; P, posterior; D, dorsal; V, ventral; CA1, cornu ammonis 1; CA2, cornu ammonis 2; CA3, cornu ammonis 3; DG, dentate gyrus; Sub, subiculum; Pro, prosubiculum; PrS/PaS, presubiculum/parasubiculum; EC, entorhinal cortex; HATA, hippocampal amygdala transition area; LGN, lateral geniculate nucleus; Fim, fimbria*.

Diagnoses of PDD were determined clinically by the donors’ physicians. All cases with PDD presented with clinical symptoms of dementia at the time of death and a neuropathological diagnosis of Parkinson disease was confirmed at post-mortem examination. Fresh-frozen tissue samples were randomly numbered by a secondary investigator (V.C.) prior to experimentation to blind primary investigators (K.J.M.) as to case diagnoses. Ethics approval was obtained from the University of Sydney Human Research Ethics Committee (approval number 2017/394). Demographic and clinical information for all cases is detailed in **Table 1**.

**Table 1.**
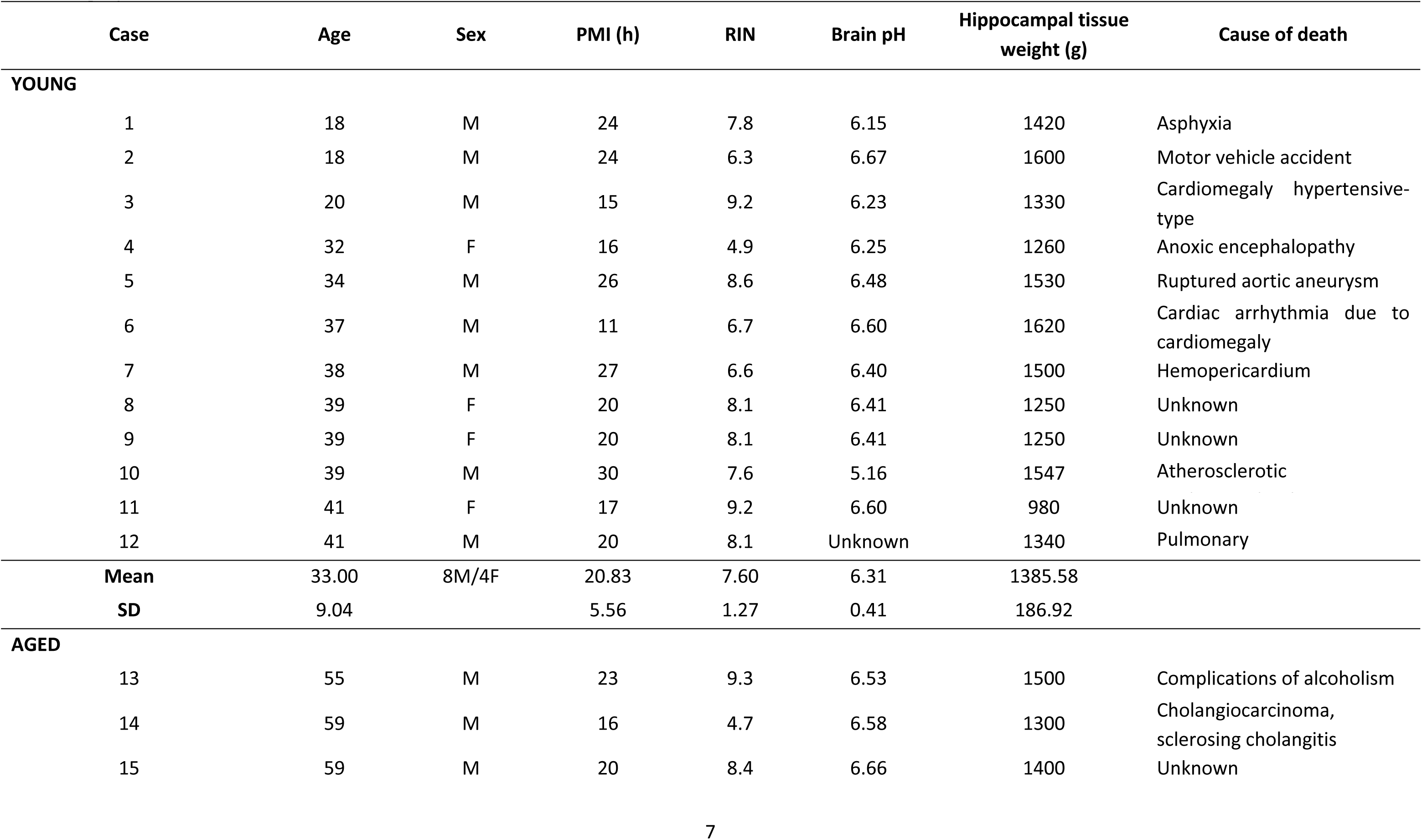
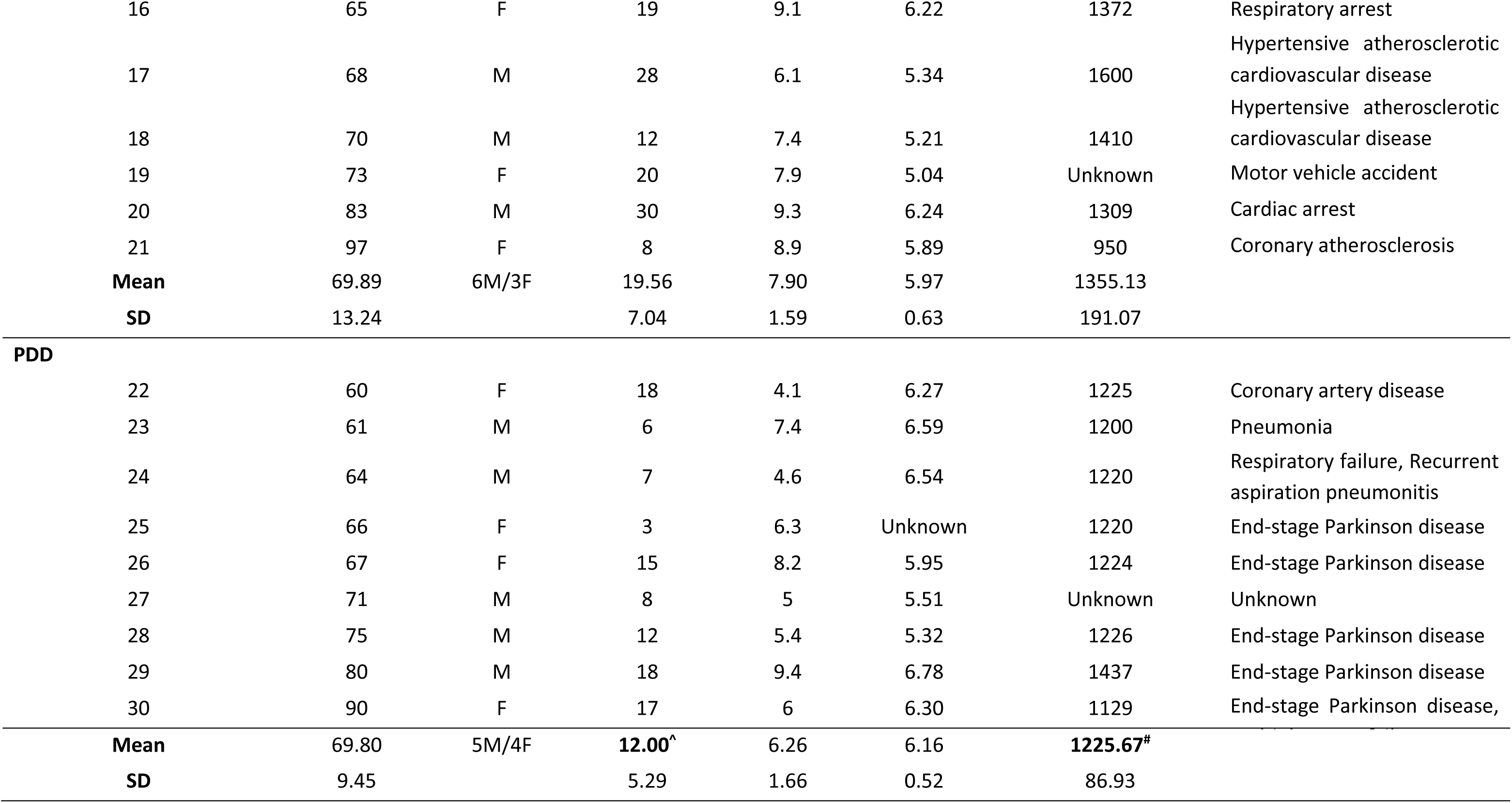
Demographic information for all cases included in the experimental cohort. ^^^PDD PMI was significantly lower than young controls (*p* = 0.01) and aged control cases (*p* = 0.05). ^#^Hippocampal tissue weight in PDD cases was significantly lower than in young controls (*p* = 0.027). Abbreviations: PMI, post-mortem interval; RIN, RNA integrity number.

### Quantification of neurogenesis-related gene expression

Neurogenesis markers were quantified at the gene transcript level. First, the central aim of this study required regional measures of neurogenesis marker expression to be paired with regional monoamine measures in the same donor, and monoamines and their metabolites are not reliably recoverable from fixed tissue. Second, immunohistochemical estimates of adult human neurogenesis are strongly influenced by post-mortem interval, fixation and tissue processing, as well as by the limited specificity of available cell type markers. Third, transcript measurement allows markers spanning successive stages of the neurogenic lineage to be assayed simultaneously from the limited tissue available when a single hippocampus is subdivided into three regions. We note that mRNA abundance does not index cell number, and the measures reported here are therefore interpreted as transcriptional signatures of the neurogenic lineage rather than as counts of newly generated neurons.

### RNA extraction

Tissue samples of 50-100 mg were pestle-homogenised in 1 mL TRIzol reagent (Thermo Fisher Scientific, Massachusetts, USA) and incubated at room temperature for 5 min to allow dissociation of the nucleotide complex. 200 µL of chloroform was added, vortexed vigorously and incubated for 3 min before centrifuging for 15 min at 12,000× *g* (4°C) to initiate phase separation. The upper aqueous phase was transferred into new centrifuge tubes, and RNA was precipitated by adding 500 µL isopropyl alcohol, incubating for 10 min at room temperature and centrifuging for 10 min at 12,000× *g* (4°C). The resulting RNA pellet was washed 3x in 1 mL 75% ethanol and re-pelleted in a centrifuge for 5 min at 7,500× *g* (4 °C) after each wash. The RNA pellet was air-dried for 5-10 min, dissolved in 50 µL RNase-free water and stored at −80°C until further use. Optical density was read at 260 nm and 280 nm using a NanoDrop One spectrophotometer (Thermo Fisher Scientific, USA) to determine RNA concentration and purity. The A260/A280 ratio of all samples was between 1.9-2.0, suggesting little to no protein or DNA contamination.

### cDNA synthesis

Complementary DNA was synthesised using a SuperScript® III First-Strand Synthesis kit (Invitrogen, California, USA) according to the manufacturer’s protocol. Briefly, 2 µg total RNA was combined with random hexamers and free nucleotides, incubated for 5 min at 65 °C and placed immediately on ice for 1 min. 10 µL cDNA synthesis mix containing 10x reaction buffer, 25mM MgCl_2_, 0.1 M Dithiothreitol, RNaseOUT™ RNase inhibitor and SuperScript® III reverse transcriptase was added to each RNA sample. cDNA synthesis was initiated at 50°C for 50 min, followed by reaction termination at 85 °C for 5 min. Samples were diluted 5× with RNase-free water to prevent the remaining reaction mix from inhibiting follow-on polymerase chain reaction amplification. Eight randomly chosen samples underwent an additional round of cDNA synthesis. These samples were pooled and serially diluted (3×) to create a standard curve to measure relative gene expression. Samples were stored at −20°C until further use.

### Quantitative reverse transcriptase polymerase chain reaction (qRT-PCR)

Relative levels of mRNA expression were measured using TaqMan® Gene Expression Assays (Applied Biosystems, California, USA; **Supplementary Table 1**). Exon-spanning probes for the genes glial fibrillary acidic protein isoform delta (*GFAPδ;* a marker of stem cells), *MKI67* (Ki67; a marker of cell proliferation), *EOMES* (eomesodermin; a marker of neuronal progenitors), *DCX* (doublecortin; a marker of immature neurons) and *GFAP* (glial fibrillary acidic protein isoform α; a marker for astrocytes and stem cells) were chosen to ensure appropriate capture of mature mRNA expression. These genes were chosen to accurately reflect the multiple stages of neurogenesis (Mathews et al., 2017). The housekeeping genes ubiquitin C, importin 8, and TATA-box binding protein were used for normalization to calculate relative mRNA expression levels.

### Quantification of hippocampal dopamine, serotonin and metabolites

Reversed phase high-performance liquid chromatography (RP-HPLC) was used to quantify hippocampal levels of DA and its metabolites, homovanillic acid (HVA) and 3,4-dihydroxyphenylacetic acid (DOPAC), as well as 5-HT and its metabolite 5-hydroxyindoleacetic acid (5-HIAA). Hippocampal tissue samples were sonicated (Branson Sonifier, Connecticut, USA) on ice in 10x volume of ice-cold homogenisation buffer (150 mM phosphoric acid and 500 µM diethylenetriaminepentaacetic acid). Homogenates were centrifuged for 25 mins at 18,000*x g* (4°C) to pellet cellular debris. An aliquot of supernatant was removed for total protein quantification, and the remaining supernatant was filtered through Amicon® 3 kDa cut-off spin filters (Merck Millipore, USA) in a centrifuge for 90 min at 14,000× *g* (4°C). Known volumes of filtrate were lyophilised overnight at room temperature and concentrated by adding half of the original volume of homogenisation buffer to the dried samples. Samples were stored at −80° until further HPLC analysis. Total protein content of the homogenised samples’ aliquots was measured using a Pierce™ BCA assay according to the manufacturer’s protocol.

The HPLC system used consisted of a Shimadzu Prominence LC-20AD pump module (Shimadzu Corporation, Kyoto, Japan) fitted with a 150 x 4.6 mm, 5 µm diameter, reverse-phase C-18 column (Phoenemenex, California, USA) maintained at 40°C. Mobile phase contained 87% (v/v) 0.01 M sodium phosphate monobasic buffer and 13% (v/v) methanol with 0.1 mM ethylenediaminetetraacetic acid (EDTA), 0.65 mM 1-octenesulfonic acid and 0.5 mM triethylamine (pH 2.81). Mobile phase was delivered at 1 mL/min, through a 0.22 µm vacuum disc filter. Samples were measured against a standard curve of known concentration created using commercially prepared DA, DOPAC, HVA, 5-HT and 5-HIAA (Sigma-Aldrich, Missouri, USA).

Twenty microlitre sample injections were performed in duplicate with a Shimadzu Prominence SIL-20A autosampler (Shimadzu Corporation, Japan) with electrochemical detection by an electrochemical detector (Antec-Leyden, Zoeterwoude, Netherlands) with a glassy carbon electrode maintained at +0.82 V versus a silver-silver chloride reference electrode. Chromatogram analysis was completed using Shimadzu LabSolutions software v5.57 (Shimadzu Corporation, Japan). Data was expressed in pg of the compound of interest per mg total protein. Data points appearing above or below the levels defined by the standard curve were removed before analysis.

### Statistical analysis

A Kruskal-Wallis H test was used to determine any significant differences in post-mortem interval (PMI), RNA integrity number (RIN), brain pH and brain weight between young, aged and PDD cases. A Mann-Whitney U test was used to confirm age matching between the aged and PDD groups.

Two-way ANOVA was performed with diagnostic group (young, aged, PDD) and hippocampal region (head, body, tail) as independent factors. Normality was assessed using the Shapiro-Wilk test and homogeneity of variance using Levene’s test. When assumptions were met, parametric one-way ANOVA was performed for each hippocampal region separately, followed by Tukey’s HSD *post hoc* test for pairwise comparisons. For data violating normality or homogeneity assumptions, Kruskal-Wallis H test was used, followed by Dunn’s *post hoc* test with Holm-Bonferroni adjustment for multiple comparisons.

A hierarchical linear regression was used to examine a range of predictors for *MKI67* and *DCX*, to determine which factors best predict neurogenesis gene expression. For both genes, a three-block regression model was designed. The first block investigated the predictive effect of PMI and brain weight, both of which were found to be significantly lower in the PDD group. The second level included gene expression level of the genes preceding the predicted gene; GFAPδ was included in the MKI67 model and GFAPδ, MKI67 and EOMES were included in the DCX model. The last block included the DA metabolite DOPAC and serotonin metabolite 5-HIAA. An additional two-block model was created to examine the predictive effect of the DA metabolite HVA and serotonin on DCX expression, when controlling for expression of GFAPδ, MKI67 and EOMES. Regression coefficients (β) with 95% confidence intervals are reported for the final block of each model. Pearson’s correlation coefficients were calculated to examine unadjusted associations between neurotransmitter levels and neurogenesis markers, both overall (pooling all subjects) and stratified by diagnostic group or hippocampal region. Significance level for all analyses was set at *p* = 0.05. All statistical tests and graphs were performed using GraphPad Prism 11.0.0 or Python. Graphs were created in GraphPad Prism 11 (GraphPad Software Inc., California, USA).

## RESULTS

### Case demographics

The cohort comprised 12 young (33.0 ± 9.0 years; 8M/4F), 9 aged (69.9 ± 13.2 years; 6M/3F), and 9 PDD cases (69.8 ± 9.5 years; 5M/4F) as described in **Table 1**. Average age did not differ between aged cases and PDD cases (*U* = 45.00, *p* = 0.730). RNA integrity number and brain pH were comparable across groups, while PDD cases had significantly lower PMI (χ²(2) = 9.53, *p* = 0.009), brain weight (χ²(2) = 7.39, *p* = 0.025), and hippocampal tissue weight (*p* = 0.027) than young and aged cases. PMI and brain weight were included as covariates in regression analyses to account for potential confounding (**Supplementary Tables 2, 3**).

### Hippocampal neurogenesis is differentially altered by ageing and PDD across the longitudinal axis

We investigated the expression of five genes that capture successive stages of hippocampal neurogenesis, from stem and progenitor cells through to neuronal maturation and integration (**Fig. 2a**). *GFAP* and its isoform *GFAPδ* are enriched in stem cells and astrocytes, *MKI67* marks actively dividing cells, *EOMES* labels intermediate neuronal progenitors, and *DCX* indexes immature neurons. Gene expression levels for these markers were compared between young, aged, and PDD individuals along the longitudinal hippocampal axis.

**Figure 2.**
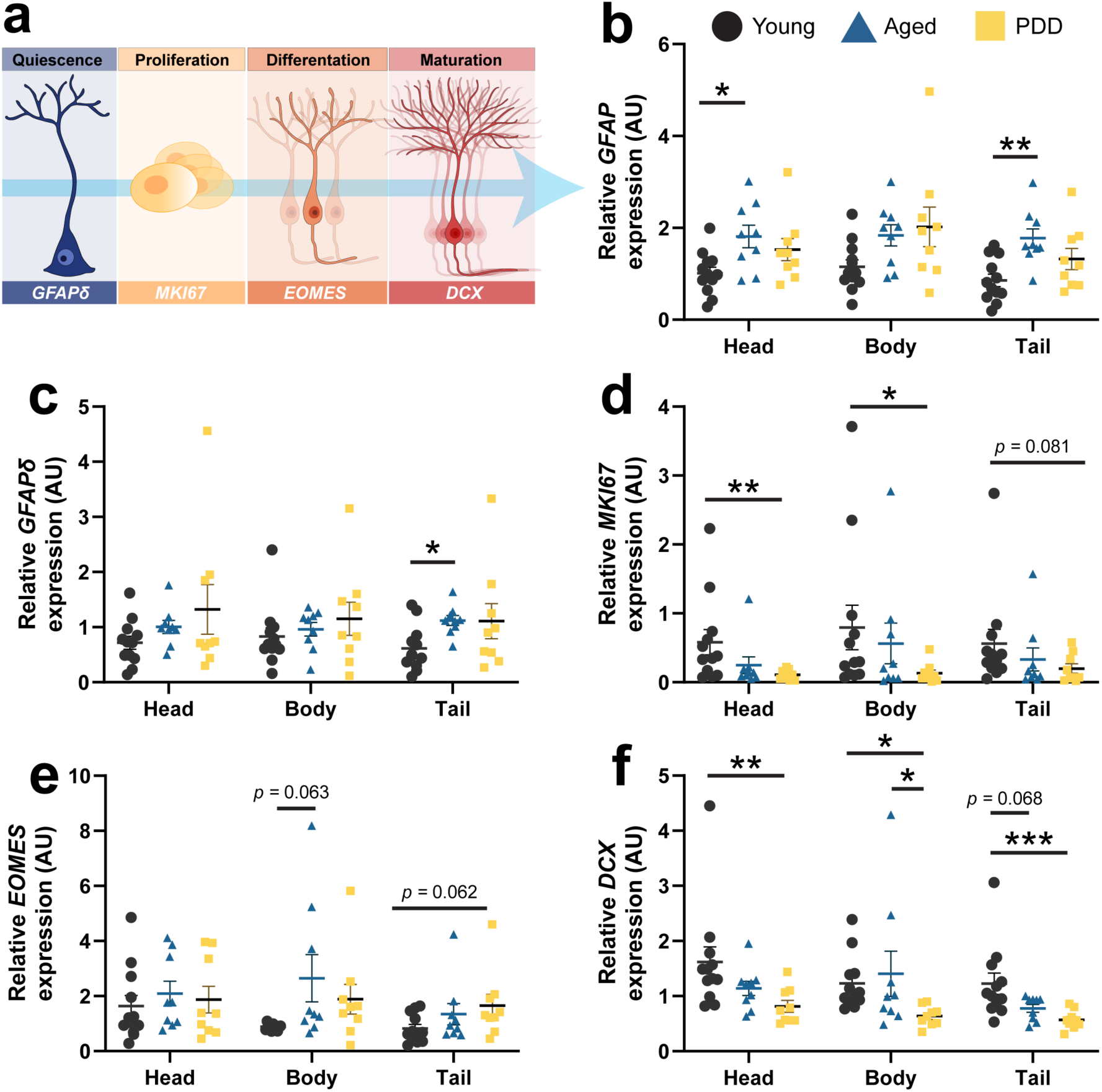
Expression of neurogenesis-associated transcripts across the hippocampal axis in young and aged controls and in PDD. Relative expression of neurogenesis-associated genes was quantified by RT-qPCR in the hippocampal head, body, and tail of young and aged controls, and in PDD. **(a)** Stages of adult hippocampal neurogenesis examined in this study, from quiescent stem cells through proliferation, differentiation, and neuronal maturation, with the corresponding marker gene indicated at each stage. **(b)** *GFAP* expression was elevated in aged controls relative to young controls in the head (*p* = 0.019) and tail (*p* = 0.003) (Group: *F*₍₂,₈₁₎ = 14.38, *p* < 0.001). **(c)** *GFAPδ* expression was elevated in aged controls compared with young controls in the tail (*p* = 0.031), with intermediate expression levels observed in PDD (Group: *F*₍₂,₈₁₎ = 4.68, *p* = 0.012). **(d)** *MKI67* expression was reduced in PDD compared with young control individuals in the head (*p* = 0.008) and tail (*p* = 0.081) (Group: *F*₍₂,₈₁₎ = 10.86, *p* < 0.001). **(e)** *EOMES* expression showed trends towards elevated levels in aged controls relative to young controls in the body (*p* = 0.063) and in PDD, relative to young controls in the tail (*p* = 0.062) (Group: *F*₍₂,₈₀₎ = 4.80, *p* = 0.011). **(f)** *DCX* expression was significantly reduced in PDD compared with young controls in the head (*p* = 0.003), body (*p* = 0.012), and tail (*p* < 0.001), with aged controls showing intermediate levels that differed from young controls in the tail (*p* = 0.068) (Group: *F*₍₂,₈₁₎ = 19.59, *p* < 0.001; Region: *F*₍₂,₈₁₎ = 4.45, *p* = 0.015). No significant interaction effects were found for the investigated gene transcripts. Brackets with asterisks denote significant pairwise comparisons from one-way ANOVAs followed by Tukey’s HSD *post hoc* or Kruskal-Wallis followed by Dunn’s *post hoc* tests. \**p* < 0.05, \*\**p* < 0.01, \*\*\**p* < 0.001. Data are shown as individual values with group means ± SEM. *n* = 9–12/group.

Expression of the astrocytic and stem cell-associated marker *GFAP* was significantly elevated by up to 51% in aged controls, compared with young controls, in the hippocampal head (*p* = 0.019) and tail (*p* = 0.003; **Fig. 2b**), but not in the hippocampal body. The stem cell-associated isoform *GFAPδ* exhibited a similar pattern, with a prominent increase in the hippocampal tail of aged controls, compared with young controls (*p* = 0.031; **Fig. 2c**), suggesting the stem cell pool is preserved or even expanded in this region in the aged hippocampus. Despite the apparent preservation of the upstream stem cell environment, expression of the proliferation marker *MKI67* was markedly reduced in PDD compared with young controls in the hippocampal head (*p* = 0.008) and tail (*p* = 0.081; **Fig. 2d**). As MK167 expression in all three hippocampal regions was not significantly reduced in aged controls compared to young controls, the decline in actively dividing precursors in PDD may primarily be driven by disease rather than age (Boldrini et al., 2018; Mathews et al., 2017).

*EOMES* expression was marginally increased in aged controls relative to young controls in the hippocampal body (*p* = 0.063), and in PDD, relative to young controls in the tail (*p* = 0.062; **Fig. 2e**). Consistent with this, the most downstream marker in the neurogenic lineage, *DCX*, exhibited the most pronounced disease-related perturbations. *DCX* expression was markedly reduced in PDD relative to young controls across the entire hippocampal axis, with 50–54% reductions in the head (*p* = 0.003), body (*p* = 0.012) and tail (*p* < 0.001; **Fig. 2f**) respectively. In the hippocampal body, *DCX* expression was also reduced in PDD compared to age-matched controls (*p* = 0.035).

### PDD is associated with reduced DOPAC and elevated 5-HIAA levels in the hippocampus

Next, to assess age- and disease-associated changes in hippocampal monoamines, we quantified levels of dopamine and serotonin and their metabolites using HPLC. Dopamine levels were reduced by 47% in the hippocampal head in PDD relative to young controls, although this did not reach statistical significance (*p* = 0.075; **Fig. 3a**). The dopamine metabolite, DOPAC, was markedly reduced by 60–63% in PDD cases compared with young controls in the hippocampal head (*p* = 0.032), body (*p* = 0.027) and tail (*p* = 0.061; **Fig. 3b**). In contrast, HVA levels were unchanged between groups (**Fig. 3c**) but varied significantly along the hippocampal axis (**Supplementary Fig. 1a**). The divergence between reduced DOPAC and preserved HVA is notable, as DOPAC reflects intraneuronal dopamine metabolism whereas HVA more closely indexes extracellular dopamine metabolism. Regional HVA concentration was highest in the hippocampal head and declined by 54–59% in the body (*p* = 0.00047) and tail (*p* = 0.0068), with a similar but attenuated pattern in aged cases (**Supplementary Fig. 1a**). Dopamine turnover showed a marginal increase in PDD cases compared with the aged group in the hippocampal head, but this did not reach statistical significance (*p* = 0.059; **Fig. 3d**).

**Figure 3.**
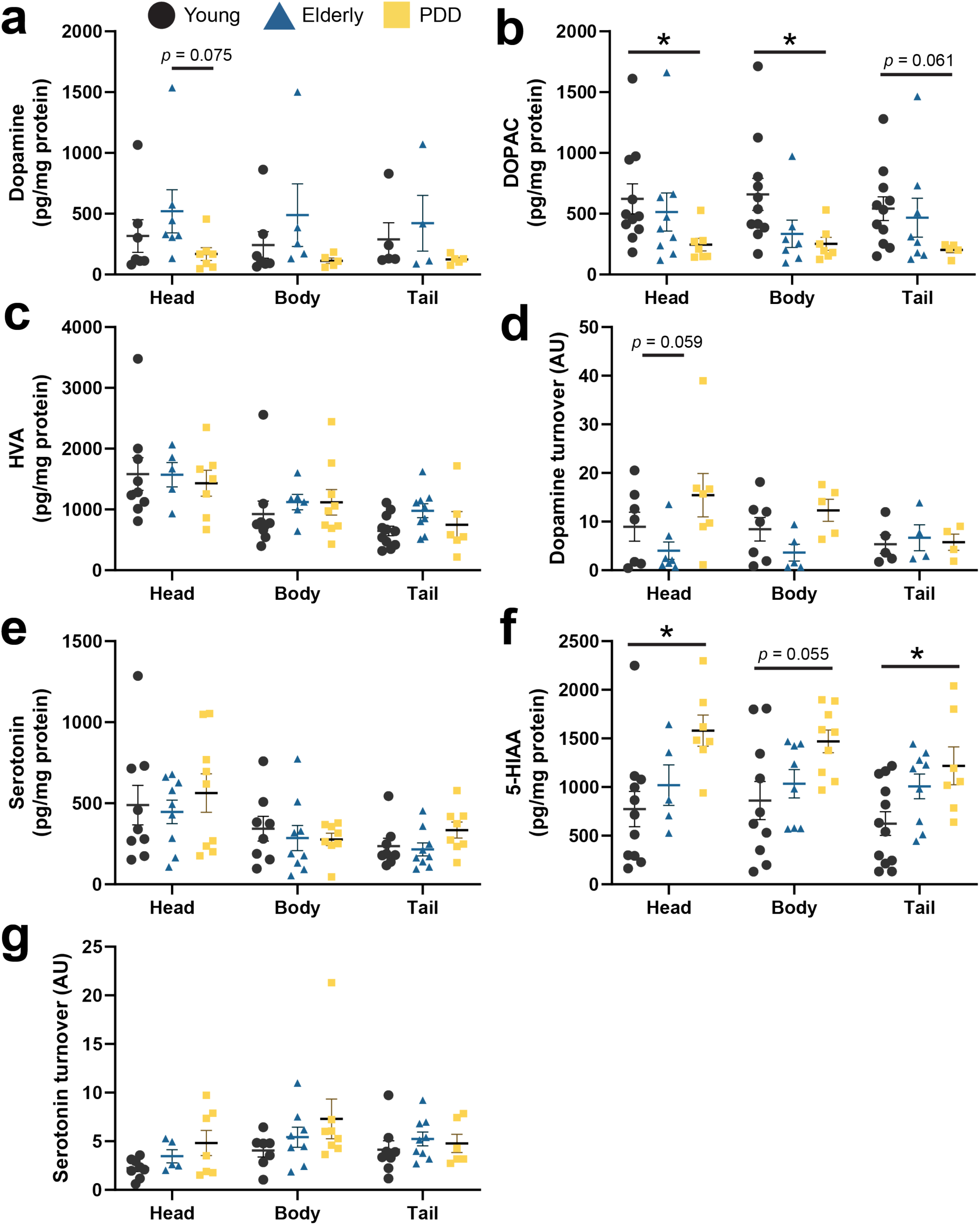
DA, 5-HT and metabolite concentrations across the hippocampal axis in young, aged and PDD donors. Tissue concentrations (pg/mg protein) of DA, 5-HT and their metabolites were quantified by HPLC in the hippocampal head, body, and tail of young (black circles), aged (blue triangles), and PDD (yellow squares) donors, with derived turnover ratios (AU). **(a)** Dopamine levels tended to be lower in PDD donors compared with young controls, primarily in the head (*p* = 0.075; Group: *F*(_2,40_) = 4.47, *p* = 0.018) **(b)** HVA concentrations did not differ between groups but showed a significant regional gradient along the hippocampal axis (Region: *F*(_2,61_) = 13.74, *p* < 0.001). **(c)** DOPAC concentrations were markedly reduced in PDD donors relative to young controls in the head (*p* = 0.032) and body (*p* = 0.027), with a trend in the tail (*p* = 0.061) (Group: *F*(_2,67_) = 6.49, *p* = 0.003). **(d)** No significant changes in dopamine turnover was observed between groups (*F*(_2,38_) = 4.99, *p* = 0.012) **(e)** Serotonin concentrations did not differ between groups but showed a significant regional gradient (*F*(_2,68_) = 7.95, *p* < 0.001) **(f)** 5-HIAA concentrations were markedly elevated in PDD relative to young controls in the head (*p* = 0.012) and tail (*p* = 0.022), with a trend in the body (*p* = 0.055) (Group: *F*(_2,69_) = 13.61, *p* < 0.001). **(g)** Serotonin turnover (5-HIAA/serotonin) showed a modest group effect (*F*(_2,56_) = 3.29, *p* = 0.045). Data are shown as individual values with group means ± SEM. Brackets with asterisks denote significant pairwise comparisons from one-way ANOVAs or Kruskal–Wallis tests. \**p* < 0.05, \*\**p* < 0.01. *n* = 4–9/group

Serotonin levels did not differ significantly across groups (**Fig. 3e**) but showed a comparable anteroposterior gradient to HVA that was most pronounced in young cases (*p* = 0.049; **Supplementary Fig. 1b**). The serotonin metabolite 5-HIAA was markedly elevated by 71–104% in the head (*p* = 0.012), body (*p* = 0.055) and tail (*p* = 0.022) in PDD compared with young controls (**Fig. 3f**). Serotonin turnover showed a modest overall group effect but no significance pairwise differences were observed (**Fig. 3g**). Taken together, these findings suggest that age-associated changes in monoamine activity in the hippocampus are more pronounced in PDD.

### Dopamine and serotonin metabolites predict alterations in neurogenesis gene expression

To identify predictors of neurogenesis markers expression beyond diagnostic group, hierarchical regression models were constructed for *DCX* and *MKI67* using a three-block design (**Supplementary Tables 2, 3**). These two markers of neurogenesis were selected because they were significantly changed with ageing and/or disease state. Model 1 included post-mortem tissue characteristics (brain weight and PMI), Model 2 added neurogenesis-associated transcripts (*GFAPδ*, *MKI67*, *EOMES*), and Model 3 incorporated the monoaminergic metabolites DOPAC and 5-HIAA (**Fig. 4a**, **Supplementary Tables 2, 3**). For *DCX*, neither post-mortem tissue characteristics nor neurogenesis transcripts alone accounted for substantial variance (**Supplementary Table 2**). The inclusion of DOPAC and 5-HIAA strengthened the full model (R² = 0.331, *p* < 0.05; **Fig 4a**) with DOPAC and *GFAPδ* emerging as significant predictors of *DCX* expression (**Fig. 4b, c**). These findings suggest that increased DOPAC levels, and reduced stem cell marker expression, predict a greater abundance of immature neurons. Interestingly, these correlations were present in aged controls (DOPAC: *r* = 0.54, *p* = 0.006; *GFAPδ*: *r* = –0.59, *p* = 0.001), but not in young controls or in PDD (**Supplementary Fig. 2**).

**Figure 4.**
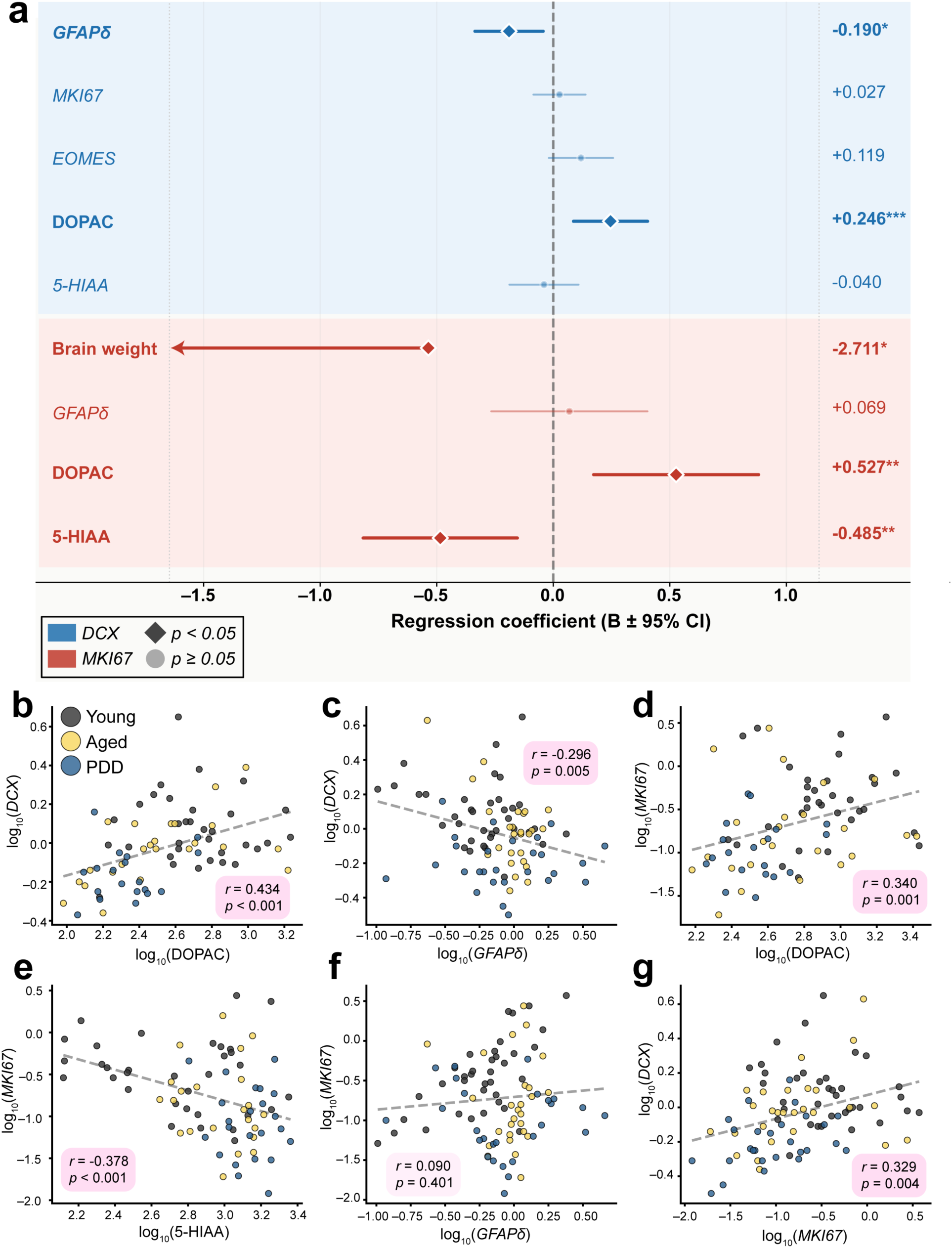
Predictors of *DCX* and *MKI67* expression identified by hierarchical linear regression modelling. **(a)** Forest plot showing standardised regression coefficients (*B* ± 95% CI) from the full model (Model 3) for *DCX* (blue) and *MKI67* (red). Dark diamonds indicate significant predictors (*p* < 0.05); light circles indicate non-significant predictors. *DCX* was independently predicted by *GFAPδ* (*B* = −0.190, *p* < 0.05) and DOPAC (*B* = +0.246, *p* < 0.001); *MKI67* was predicted by brain weight (*B* = −2.711, *p* < 0.05), DOPAC (*B* = +0.527, *p* < 0.01), and 5-HIAA (*B* = −0.485, *p* < 0.01). Full model statistics for each hierarchical step are reported in Supplementary Tables 2 and 3. **(b)** DOPAC was positively correlated with *DCX* across the full cohort (*r* = 0.434, *p* < 0.001); within-group analysis revealed this was driven by aged controls (*r* = 0.54, *p* = 0.006), with no significant correlations present for young controls or in PDD. **(c)** *GFAPδ* was inversely correlated with *DCX* (*r* = −0.296, *p* = 0.005), most strongly in aged donors (*r* = −0.59, *p* = 0.001). **(d)** DOPAC was positively correlated with *MKI67* (*r* = 0.340, *p* = 0.001). **(e)** 5-HIAA was inversely correlated with *MKI67* (*r* = −0.378, *p* < 0.001). **(f)** *GFAPδ* was not correlated with *MKI67* across the full cohort (*r* = 0.090, *p* = 0.401). There was, however, a strong positive correlation between these factors in young donors (*r* = 0.69, *p* < 0.001) that was absent in aged controls (*r* = −0.04) and in PDD (*r* = −0.07). **(g)** *DCX* and *MKI67* were positively correlated in the whole cohort (*r* = 0.329, *p* = 0.004), consistent with co-variation of proliferative activity and immature neuron abundance. Scatter points are coloured by diagnostic group (black, young; yellow, aged; blue, PDD). Overall Pearson correlation coefficients and *p*-values are shown on each panel. \**p* < 0.05, \*\**p* < 0.01, \*\*\**p* < 0.005.

Similar to *DCX*, the first two blocks of the model did not account for the significant variance observed in *MKI67* (**Supplementary Table 3**). The addition of DOPAC and 5-HIAA to the full model produced a substantial increase in the strength of these relationships (R² = 0.261, *p* < 0.001), indicating that monoaminergic metabolites are a strong predictor of proliferative activity. This is supported by independently correlations between both DOPAC and 5-HIAA with *MKI67* (both *p* ≤ 0.001; **Fig. 4d, e**). Brain weight was also negatively associated with *MKI67* expression in the full model, suggesting donors with greater brain atrophy tend to have reduced neuronal proliferation. Unlike *DCX*, *GFAPδ* did not predict *MKI67* across the full cohort (**Fig. 4f**), but we found a strong positive correlation between these factors in young donors (*r* = 0.69, *p* < 0.001), a relationship that was absent in the elderly and PDD groups. This suggests that the functional coupling between stem cell marker expression and active proliferation, which is intact in the young hippocampus, is disrupted in both normal ageing and neurodegeneration. *DCX* and *MKI67* were positively correlated across the cohort (*r* = 0.329, *p* = 0.004; **Fig. 4g**), consistent with the expectation that proliferative activity is associated with immature neuron abundance.

## DISCUSSION

In the present study, we demonstrate that transcriptional signatures associated with the maturation of adult hippocampal neurons are diminished with age along the hippocampal longitudinal axis, but are most perturbed in individuals with a neuropathological disease affecting memory and cognition. These changes co-occur with region-specific alterations in dopaminergic and serotonergic metabolism that are predominantly disease-driven. Hierarchical regression modelling revealed that these monoaminergic metabolites are significant predictors of proliferation and maturation markers, implicating monoaminergic dysregulation as a potential contributor to the reduced neurogenesis phenotype in ageing and PDD.

Studies using carefully phenotyped post-mortem tissue indicate that adult hippocampal neurogenesis follows a staged pattern across the human lifespan (Boldrini et al., 2018; Mathews et al., 2017). We and others have previously shown that stem cells remain relatively stable across the lifespan while the proliferation and maturation of neurons declines with age (Knoth et al., 2010; Mathews et al., 2017; Moreno-Jiménez et al., 2021; Spalding et al., 2013). By mapping the expression of neurogenesis markers along the hippocampal axis, we found an age-associated decoupling between the stem cell-like marker *GFAPδ* and *DCX* expression in the tail of the hippocampus. The observation of these regionally confined changes indicates that bulk analysis of the whole hippocampus does not capture region-specific changes in neurogenesis marker expression. Importantly, this pattern suggests that ageing preferentially shifts the hippocampal tail towards an accumulation of quiescent stem cell-like and astroglial populations without a region-specific depletion of the proliferating or maturing neuronal compartment, a distinction that becomes important when contrasting the ageing phenotype with the more severe depletion seen in PDD.

In our cohort, PDD cases show pronounced reductions in both *MKI67* and *DCX* expression beyond those observed in aged individuals, indicating that the disease process can exacerbate nascent age-associated vulnerability to impair the proliferation and maturation of granule neuronal precursors. The strongest reductions in PDD are concentrated in the hippocampal tail, the region most critical for spatial and episodic processing, suggesting that disease-related neurogenic impairment may contribute directly to the pattern of cognitive deficits characteristic of PDD (Bouchard et al., 2008; Kehagia, Barker, & Robbins, 2010). Critically, the reduction in cell maturation in PDD is not accompanied by further expansion of stem cell-like or astroglial populations beyond that already established by ageing, indicating that the disease selectively depletes the proliferating and maturing compartments from a niche that ageing has already shifted towards quiescence. Recent evidence reported a significant reduction in DCX^+^ progenitor cells and their nuclear size in the subgranular zone in Parkinson disease, indicating impaired maturation and integration of immature neurons in cognitively intact patients (Plácido, Koss, Outeiro, & Brocardo, 2024). These findings complement other studies indicating that perturbations to hippocampal neurogenesis in Parkinson disease may contribute to the onset of non-motor symptoms such as depression and early cognitive vulnerability (Alonso-Moreno, Gallardo-Caballero, Prádanos-Senén, & Llorens-Martín, 2025; Lim, Bang, & Choi, 2018). In Alzheimer’s disease, several studies report a marked loss of proliferating progenitors and doublecortin-positive immature neurons in the dentate gyrus, together with abnormal morphology of the remaining immature cells, with the severity of this impairment associated with both Braak stage and cognitive decline (Knoth et al., 2010; Moreno-Jiménez et al., 2021; Salta et al., 2023; Tobin et al., 2019). Similar reductions or distortions in neurogenesis markers have also been described in dementia with Lewy bodies and mixed Lewy body pathologies, where progenitor populations and the expression of neurogenic markers are altered in the hippocampus (Johnson et al., 2011; Terreros-Roncal et al., 2021). These observations indicate that adult hippocampal neurogenesis is particularly vulnerable to neurodegenerative pathology across aetiologies and may be affected early in the transition from prodromal stages to clinically overt dementia.

To relate these transcriptional changes to the monoaminergic environment, we examined how dopamine and serotonin levels and metabolism changed across the hippocampal axis. The monoaminergic alterations in our cohort are predominantly a feature of disease rather than normal ageing. 5-HIAA was significantly increased in PDD relative to young adults in the hippocampal head and tail, while DOPAC was markedly reduced in the PDD head and body but relatively preserved in aged cases. Our dopaminergic findings align with studies demonstrating loss of dopamine transporter and altered dopamine homeostasis in hippocampus and limbic regions in PDD, indicative of reduced dopaminergic innervation accompanied by compensatory changes in turnover (Buddhala et al., 2015; Pavese, Metta, Bose, Chaudhuri, & Brooks, 2010). Remarkably, DOPAC and HVA revealed strikingly divergent patterns. DOPAC reflects intraneuronal dopamine metabolism, whereas HVA more closely indexes extracellular dopamine metabolism, suggesting that the reduction in DOPAC levels in PDD is consistent with loss of dopaminergic terminals rather than altered extracellular metabolism. The hippocampal head, which receives dense dopaminergic input from the ventral tegmental area (Tsetsenis, Broussard, & Dani, 2023b), showed the strongest dopaminergic alterations. This anteroposterior gradient is consistent with the topographic organisation of midbrain dopaminergic projections to the ventral hippocampus in rodents, considered analogous to the hippocampal head in humans (Tsetsenis et al., 2023b). This pattern is compatible with compensatory increases in dopamine turnover in surviving terminals, analogous to well-described adaptations in the nigrostriatal system (Sossi et al., 2002).

Our hierarchical regression model revealed that higher levels of the dopamine metabolite DOPAC were associated with higher expression of *MKI67* and *DCX*, whereas higher levels of the serotonin metabolite 5-HIAA were linked to lower *MKI67* expression. Because DOPAC and 5-HIAA reflect dopamine and serotonin turnover, respectively, these associations support a model in which regions with greater effective dopaminergic tone retain more proliferating precursors and immature neurons, whereas increased serotonin metabolism is associated with reduced proliferative capacity. This accords with experimental work demonstrating that dopamine promotes adult hippocampal neurogenesis, largely via D1-like receptor signalling, and that dopaminergic depletion reduces precursor proliferation in the dentate gyrus (Höglinger et al., 2004; Salvi et al., 2016; Takamura et al., 2014). Serotonin is likewise a positive regulator of adult hippocampal neurogenesis, with enhanced serotonergic transmission increasing cell proliferation and survival of newborn granule neurons (Banasr, Hery, Printemps, & Daszuta, 2004; Benninghoff et al., 2010; Gould & Tanapat, 1999; Segi-Nishida, 2017). Our observed negative association between 5-HIAA and the proliferation marker *MKI67* does not contradict these findings but rather suggests that 5-HIAA likely reflects compensatory turnover within a degenerating serotonergic system, instead of enhanced synaptic serotonin availability. Given that dopaminergic perturbations are selective to the region most strongly coupled to limbic and affective networks, these neurochemical changes may also contribute to the neuropsychiatric symptoms that frequently precede cognitive decline in PDD (Aarsland et al., 2017). Together, the relationship between monoamines and neurogenic transcripts provides mechanistic insights into the stage-specific erosion of neurogenic lineage observed with ageing and neurodegeneration.

In conclusion, our data provide convergent molecular evidence that adult hippocampal neurogenesis undergoes a two-stage disruption, with modest regionally patterned remodelling towards quiescence during normal ageing that is substantially accelerated in PDD. Ageing preferentially accumulates quiescent stem cell-like and astroglial populations without substantially reducing neurogenic transcript expression, whereas PDD, acting on this already-remodelled niche, produces the strongest reductions in neurogenic transcripts, particularly *DCX* in the hippocampal tail, a region critical for high-fidelity episodic and spatial memory. By integrating transcriptional and neurochemical profiling in well-characterised human post-mortem tissue, our study identifies disease-driven monoaminergic dysregulation as a key mechanistic link to compromised hippocampal neurogenesis in PDD.

## Acknowledgments

This study was funded by the Sir Zelman Cowen Universities Fund, donated by family of the late Mr Leslie Rich. Human tissue was received from the University of Maryland Brain and Tissue Bank, a biorepository of the NIH NeuroBioBank.

## Data availability statement

All associated data from this manuscript are available from the corresponding author upon reasonable request.

## SUPPLEMENTARY INFORMATION

### SUPPLEMENTARY TABLES

**Supplementary Table 1.**
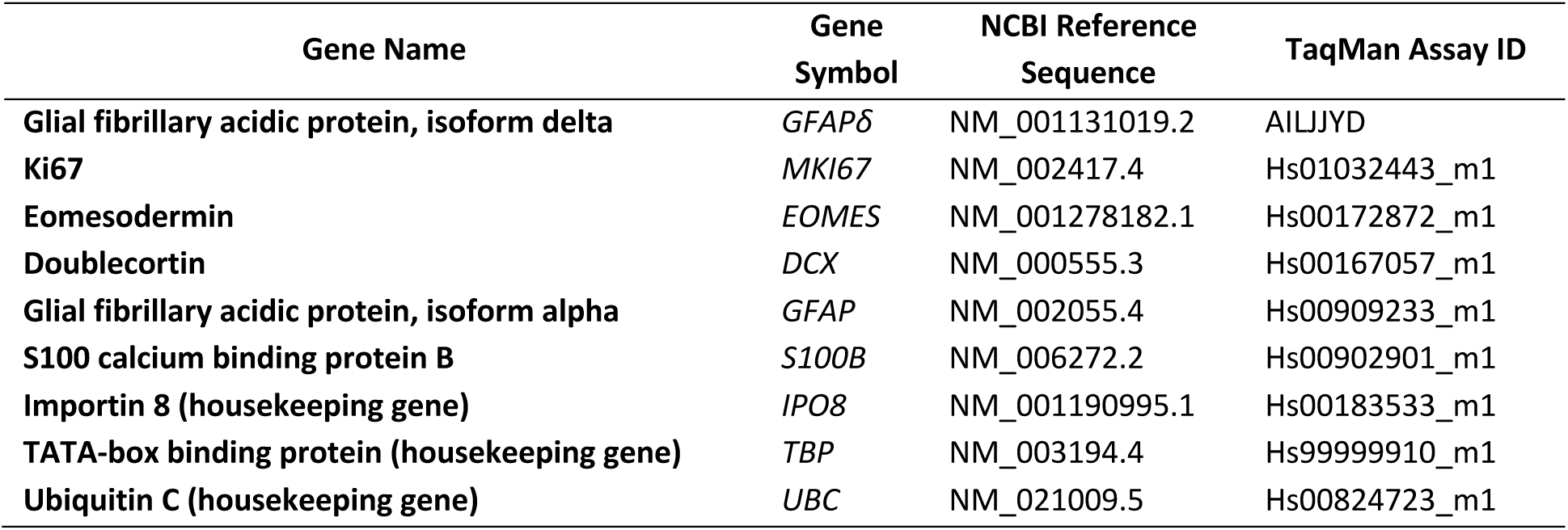
TaqMan probes used for gene expression analysis.

**Supplementary Table 2.** Hierarchical linear mixed model to determine group- and region-specific demographic, genetic and metabolic predictors of *DCX* expression.

| ***DCX* – Group-associated predictors** |  |  |  |  |  |  |
|  | *Model 1* |  | *Model 2* |  | *Model 3* |  |
| Independent variable | B | SD | B | SD | B | SD |
|---|---|---|---|---|---|---|
| PMI | 0.201 | 0.126 | 0.087 | 0.128 | 0.005 | 0.123 |
| Brain weight | 0.324 | 0.493 | 0.455 | 0.477 | 0.004 | 0.487 |
| *GFAPδ* |  |  | **-0.174\*** | 0.080 | **-0.190\*** | 0.075 |
| *MKI67* |  |  | 0.103 | 0.052 | 0.026 | 0.057 |
| *EOMES* |  |  | 0.107 | 0.075 | 0.118 | 0.070 |
| DOPAC |  |  |  |  | **0.248\*\*\*** | 0.081 |
| 5-HIAA |  |  |  |  | -0.040 | 0.076 |
| *R<sup>2</sup>* | 0.074 |  | 0.207 |  | 0.331 |  |
| *F* | 2.269 |  | **2.811\*** |  | **3.679\*\*\*** |  |
| Δ*R<sup>2</sup>* | 0.074 |  | 0.133 |  | 0.125 |  |
| Δ*F* | 2.269 |  | **3.013\*** |  | **4.847\*** |  |

**Supplementary Table 3.** Three-block hierarchical regression models to determine group-specific demographic, genetic and metabolic predictors of *MKI67* expression.

| ***MKI67* – Group-associated predictors** |  |  |  |  |  |  |
|  | *Model 1* |  | *Model 2* |  | *Model 3* |  |
| Independent variable | B | SD | B | SD | B | SD |
|---|---|---|---|---|---|---|
| PMI | 0.285 | 0.307 | 0.357 | 0.326 | 0.116 | 0.295 |
| Brain weight | -1.324 | 1.204 | -1.248 | 1.215 | **-2.709\*** | 1.109 |
| *GFAPδ* |  |  | 0.134 | 0.195 | 0.067 | 0.171 |
| DOPAC |  |  |  |  | **0.528\*\*** | 0.180 |
| 5-HIAA |  |  |  |  | **-0.484\*\*** | 0.169 |
| *R<sup>2</sup>* | 0.025 |  | 0.033 |  | 0.228 |  |
| *F* | 0.748 |  | 0.650 |  | **4.540\*\*** |  |
| Δ*R<sup>2</sup>* | 0.025 |  | 0.008 |  | 0.259 |  |
| Δ*F* | 0.748 |  | 0.469 |  | **10.065\*\*\*** |  |

\* = *p* < 0.05, \*\* = *p* < 0.01, \*\*\* = *p* < 0.005

### SUPPLEMENTARY FIGURES

**Supplementary Figure 1.**
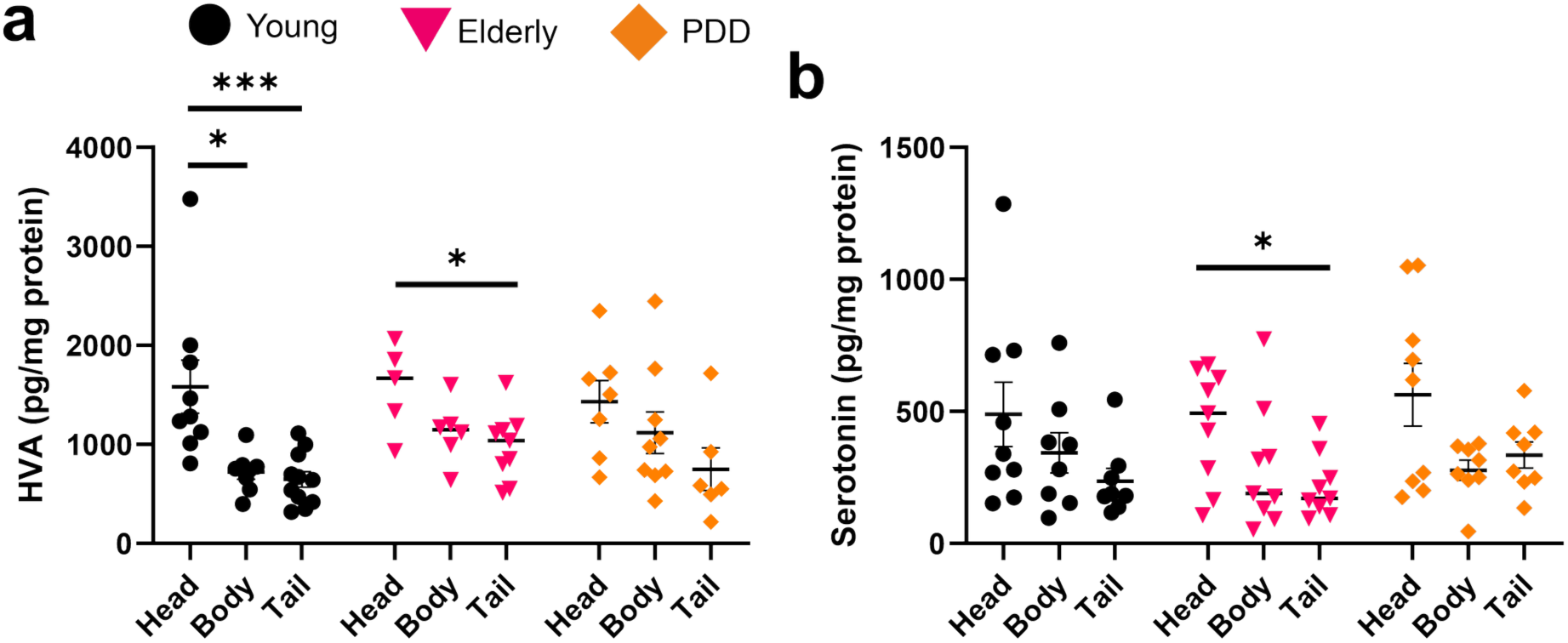
Regional distribution of HVA and serotonin across the hippocampal axis in young, aged, and PDD donors. Monoamine concentrations were quantified by HPLC in the hippocampal head, body, and tail of young and aged controls and in PDD. (**a**) HVA concentrations showed a significant anterior-to-posterior gradient (Region: F₍₂,₆₁₎ = 12.69, *p* < 0.001) with no significant group effect (Group: F₍₂,₆₁₎ = 1.34, *p* = 0.269). In young donors, HVA was significantly higher in the head than the body (*p* = 0.007) and tail (*p* < 0.001). In aged donors, HVA was higher in the head than the tail (*p* = 0.025). No significant regional differences were observed in PDD. (**b**) Serotonin concentrations likewise showed a significant regional gradient (Region: F₍₂,₆₈₎ = 7.84, *p* < 0.001) with no significant group effect (Group: F₍₂,₆₈₎ = 0.68, *p* = 0.508). Aged donors showed significantly higher serotonin in the head than the tail (*p* = 0.049). No significant interaction effects were found for either analyte. Brackets with asterisks denote significant pairwise comparisons from one-way ANOVAs followed by Tukey’s HSD post hoc or Kruskal–Wallis followed by Dunn’s post hoc tests. \**p* < 0.05, \*\**p* < 0.01, \*\*\**p* < 0.001. Data are shown as individual values with group means ± SEM.

**Supplementary Figure 2.**
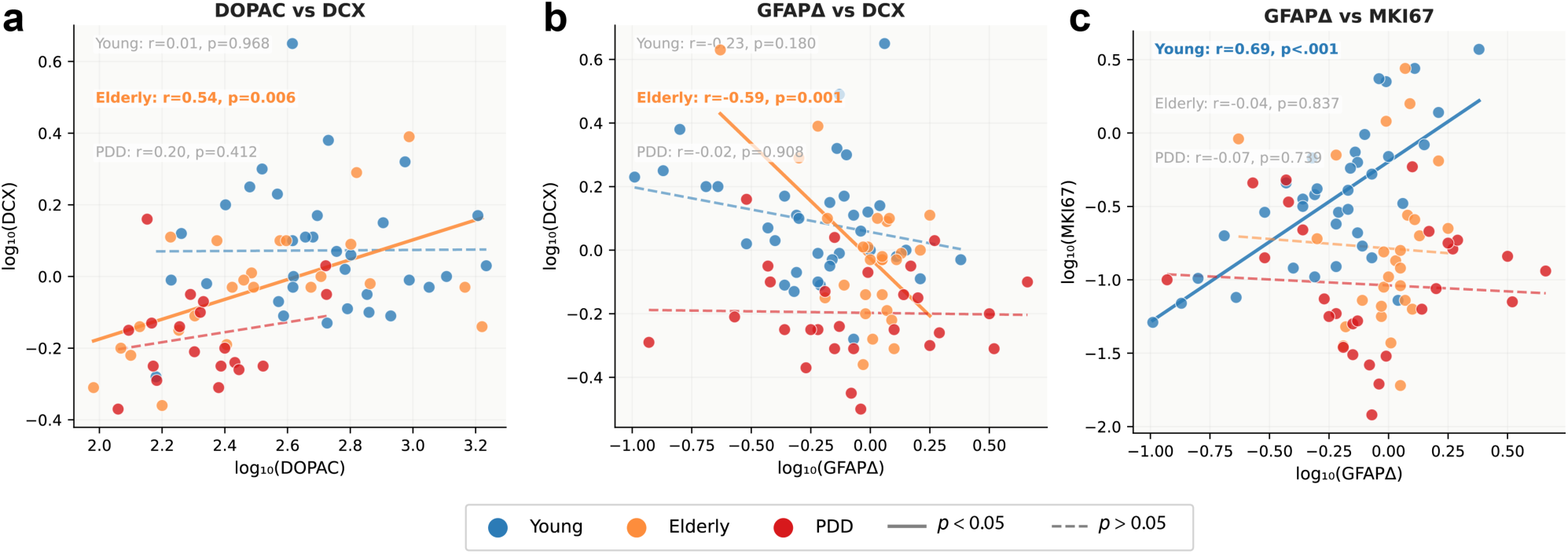
Correlations between monoaminergic and neurogenesis-associated markers. Correlations shown in main Figure 4 were stratified by diagnostic group to examine whether relationships between predictors and neurogenesis markers were consistent across young and aged controls and in PDD. (**a**) DOPAC was positively correlated with *DCX* in aged donors (*r* = 0.54, *p* = 0.006) but not in young donors (*r* = 0.01, *p* = 0.968) or in PDD (*r* = 0.20, *p* = 0.412). (**b**) *GFAPδ* was inversely correlated with DCX in aged donors (*r* = −0.59, *p* = 0.001) but not in young donors (r = −0.23, p = 0.180) or in PDD (r = −0.02, p = 0.908). (**c**) *GFAPδ* were inversely correlated with *MKI67* expression levels in young donors (*r* = −0.69, *p* < 0.001) but not in aged donors (r = 0.04, p = 0.837) or in PDD (*r* = −0.01, *p* = 0.738).

